# A Role for the Fornix in Temporal Sequence Memory

**DOI:** 10.1101/2022.08.01.498998

**Authors:** Marie-Lucie Read, Katja Umla-Runge, Andrew D. Lawrence, Alison G. Costigan, Liang-Tien Hsieh, Maxime Chamberland, Charan Ranganath, Kim S. Graham

**Affiliations:** Cardiff University Brain Research Imaging Centre (CUBRIC), School of Psychology, Cardiff University, Cardiff, United Kingdom; School of Medicine, Cardiff University, Cardiff, United Kingdom; Helen Willis Neuroscience Institute, University of California at Berkeley, California, USA; Center for Neuroscience, Department of Psychology, University of California at Davis, Davis, California, USA; Donders Institute for Brain, Cognition and Behavior, Radboud University, Nijmegen, The Netherlands; Department of Psychology, University of Edinburgh, Edinburgh, United Kingdom

**Author notes:** **Corresponding author’s email address:**. Co-first authors (Marie-Lucie Read, Katja Umla-Runge). **Ethics Approval Statement:** This research was approved by the Cardiff University School of Psychology Research Ethics Committee.

**Keywords:** episodic memory, diffusion MRI, fornix, hippocampus, time, sequence

## Abstract

Converging evidence from studies of human and nonhuman animals suggests that the hippocampus contributes to sequence learning by using temporal context to bind sequentially occurring items. The fornix is a white matter pathway containing the major input and output pathways of the hippocampus, including projections from medial septum, and to diencephalon, striatum, and prefrontal cortex. If the fornix meaningfully contributes to hippocampal function, then individual differences in fornix microstructure might predict sequence memory. Here, we tested this prediction by performing tractography in 51 healthy adults who had undertaken a sequence memory task. Microstructure properties of the fornix were compared with those of tracts connecting medial temporal lobe regions, but not predominantly the hippocampus: the Parahippocampal Cingulum bundle (PHC) (conveying retrosplenial projections to parahippocampal cortex) and the Inferior Longitudinal Fasciculus (ILF) (conveying occipital projections to perirhinal cortex). Using principal components analysis, we combined Diffusion Tensor Imaging and Neurite Orientation Dispersion and Density Imaging measures obtained from multi-shell diffusion MRI into two informative indices, the first (PC1) capturing axonal packing/myelin, the second (PC2) capturing microstructural complexity. We found a significant correlation between fornix PC2 and implicit reaction-time indices of sequence memory, indicating that greater fornix microstructural complexity is associated with better sequence memory. No such relationship was found with measures from the PHC and ILF. This study highlights the importance of the fornix in aiding memory for objects within a temporal context, potentially reflecting a role in mediating network communication within an extended hippocampal system.

## 1.0. Introduction

The hippocampus is widely known to make a critical contribution to episodic memory. Considerable evidence suggests that the key contribution of the hippocampus might be to build memories that associate different pieces of information occuring in a shared spatiotemporal context (Aggleton & Brown, 2005; Ekstrom & Ranganath, 2018; Hsieh et al., 2014; Long & Kahana, 2019; Opitz, 2014). For instance, hippocampal lesions in animals disrupt memory for temporal order relationships (Heuer & Bachevalier, 2013; Ranganath & Hsieh, 2016). Furthermore, hippocampal neural activity patterns, measured with functional magnetic resonance imaging (fMRI), carry information about the temporal context of items in learned sequences (Hsieh et al., 2014; Kalm et al., 2013; Williams et al., 2020; for review see Lee et al., 2020).

In particular, work by Hsieh et al. (2014) evidenced that the hippocampus uniquely carries information about the temporal order of objects in sequences, whereas other Medial Temporal Lobe (MTL) regions, the perirhinal cortex and parahippocampal cortex, carry information about isolated object and temporal context information, respectively. Additionally, individual differences in sequence memory (indexed by the reaction time, RT, reduction for semantic decisions on objects within consistently-ordered versus randomly-ordered learned sequences) strongly correlated with individual differences in hippocampal object-position binding bilaterally (indexed as the multi-voxel pattern similarity difference between hippocampal patterns during repetitions of the same item in consistent sequences and hippocampal patterns during repetitions of the same item in random sequences). The results indicate that the hippocampus has computational specializations that support sequence memory.

However, it is clear that episodic memory relies, not only on the hippocampus, but also on a broader network of areas, so we aimed to elucidate how structural connectivity of the hippocampus to these broader areas relates to sequence memory. According to one hypothesis, episodic memory is supported by an “extended hippocampal system”, comprising the hippocampus, anterior thalamic nucleus, mamillary bodies and medial prefrontal cortex (Aggleton, 2012; Aggleton & Brown, 1999; Gaffan, 1992; Gaffan & Hornak, 1997). Murray et al. (2017, 2018) proposed that the prominance of integrated spatiotemporal representations in the primate extended hippocampal system reflects an inheritence from oligocene anthropoids, where background scenes supported foraging choices at a distance, an ability that was enhanced over evolution, enabling hippocampal representations to support spatio-temporal attributes of episodic memory in humans (Murray et al., 2017). Regions in the extended hippocampal system are connected via the fornix, the main white matter tract entering and exiting the hippocampus (Aggleton, 2012; Aggleton & Brown, 1999; Bubb et al., 2017; Gaffan, 1992; Gaffan & Hornak, 1997). Thus, the fornix may play a critical role in bringing together elements of the extended hippocampal system in the service of episodic memory. Individual differences in microstructure of the fornix should therefore be associated with inter-individual variation in memory for objects in spatial and/or temporal context.

Indeed, human studies using diffusion-weighted Magnetic Resonance Imaging (dMRI), which can non-invasively delineate the path of major fibre pathways and evaluate their microstructure through indices such as Fractional Anisotropy (FA) (Assaf et al., 2019), have found that inter-individual differences in fornix microstructure in healthy individuals correlate with individual variation in spatial memory and scene discrimination performance (Bourbon-Teles et al., 2021; Hodgetts et al., 2015; Hodgetts et al., 2017; Postans et al., 2014; Rudebeck et al., 2009). However, it is unclear whether fornix microstructure would similarly relate to temporal memory. While lesions of the fornix in macaque monkeys are known to impair choices based on relative recency (Charles et al., 2004), the contribution of the fornix to temporal sequence memory has not been investigated in healthy humans.

We investigated fornix contributions to memory for objects in temporal context, in healthy humans, using an adaptation of the sequence memory task of Hsieh et al. (2014). As in that study, individual differences in the time taken to answer semantic questions about learned objects in consistently ordered sequences versus randomly ordered sequences (RT random - RT consistent) was used as a measure of sequence memory (referred to as ‘Sequence Memory Performance’). Larger Sequence Memory Performance scores would equate to a faster RT for consistently ordered sequences than randomly ordered sequences, indicating that temporal context memory had aided performance (note that previous work referred to this measure as “RT Enhancement”; Hsieh et al., 2014). It was hypothesized that individual differences in fornix microstructure would relate to Sequence Memory Performance.

In contrast, we investigated the roles of the Parahippocampal Cingulum bundle (PHC) and Inferior Longitudinal Fasciculus (ILF), specifically with the expectation that these would not contribute to binding of objects within temporal sequences. The PHC connects the posterior cingulate and retrosplenial cortex with the parahippocampal cortex (Bubb et al., 2017; Bubb et al., 2018), an area previously shown to hold temporal context information about sequences, but not conjoined object and temporal position information (Hsieh et al., 2014). The ILF is a major white matter tract in the ventral object processing stream, connecting occipital cortex with ventro-anterior temporal lobe, including the perirhinal cortex (Catani et al., 2003; Hodgetts et al., 2015), an area previously shown to hold object information regardless of any temporal sequence associations, but not conjoined object and temporal position information (Hsieh et al., 2014; see also Parker & Gaffan, 1996, 1997 for related evidence of a unique role for the hippocampus-fornix system, but not perirhinal or cingulate cortices, in the integration of objects and their position in space).

A multi-shell diffusion MRI protocol was applied allowing the combination of Diffusion Tensor Imaging (DTI) and Neurite Orientation Dispersion and Density Imaging (NODDI) models to assess tract microstructural properties. While DTI has been successfully applied to study inter-individual differences in cognition and tract microstructure in healthy adults (e.g., Coad et al., 2020; Hodgetts et al., 2015; Hodgetts et al., 2017; Postans et al., 2014), DTI measures lack biological specificity (Tournier et al., 2011). For example, both axon density and fibre dispersion can influence FA (Beaulieu, 2002). Being a multi-compartment model, NODDI provides metrics sensitive to axon density, Neurite Density Index (NDI), and the extent of orientational dispersion within a voxel, Orientation Dispersion Index (ODI) (Zhang et al., 2012). Although individual diffusion measures are related to somewhat different aspects of white matter microstructure, they also share information (Bells et al., 2011; De Santis et al., 2014). A dimensionality reduction framework, based on Principal Components Analysis (PCA), can take advantage of such redundancies to combine diffusion-measures into more biologically informative measures of white matter microstructure (Chamberland et al., 2019; Geeraert et al., 2020). By adopting this novel approach, we were able to combine dMRI measures into meaningful indices of white matter microstructure, and successfully examine the relationships between white matter microstructure of the fornix, ILF and PHC, and individual differences in sequence memory. Based on Hsieh et al. (2014), we predicted significant correlations between Sequence Memory Performance and fornix microstructure properties, but not between Sequence Memory Performance and ILF or PHC microstructure properties.

## 2.0. Materials and Methods

### 2.1. Participants

Fifty-one female volunteers (mean age: 20.1 years, SD 1.1, range: 19-24 years) were recruited from the Cardiff University School of Psychology participant panel (a subsample of those described in Karahan et al., 2019). The study was approved by Cardiff University’s School of Psychology Research Ethics Committee, and all volunteers gave informed consent prior to participation. Participants underwent behavioural testing followed by a diffusion MRI scan, within a 6-month period.

### 2.2. Task procedures

The sequence memory task, adapted from Hsieh et al. (2014), comprised two sessions, a *learning session*, immediately followed by a *retrieval session*. In these, participants were asked to make semantic decisions about objects, including man-made objects, animals, fruits, and vegetables, that were presented in sequences of five objects. Participants answered different semantic questions in each of the two sessions, to ensure their responses were modulated by their memory of the temporal-sequential relationships among the objects, rather than learning at the level of motor responses or of object-response associations. Both sessions included consistently ordered (henceforth, **consistent**) and randomly-ordered (henceforth, **random**) sequence types. Consistent sequences contained the same objects that always appeared in a fixed order: one of these sequences contained unique objects and another two sequences shared identical objects in serial positions 2 and 3 (but note that this is treated as one condition as in Crivelli-Decker et al., 2018). Two random sequences contained unique objects, but these were presented in a different order in every repetition. The retrieval session additionally included novel sequences, which contained novel and trial-unique objects upon every repetition (note: response RTs to novel objects is not included in this study). Examples of the sequences are shown in Figure 1.

**Figure 1.**
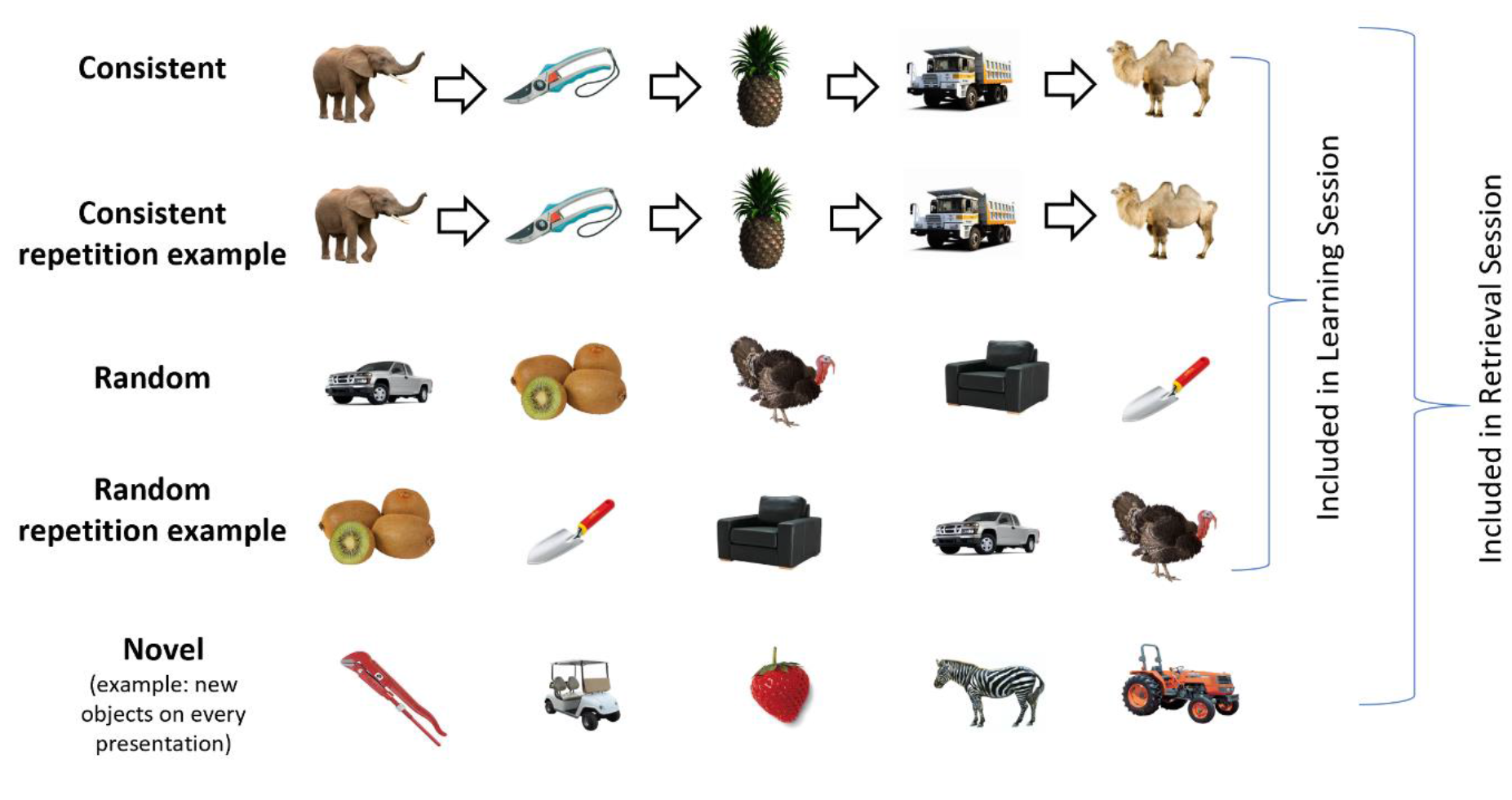
Examples of Object Sequences. Participants learned, and were tested on, a range of object sequences that were either consistently ordered (“consistent”; arrows indicate consistent order), or randomly ordered (“random”). Sequences of novel objects (“novel”) were additionally included in the retrieval stage. Repetition examples are included to illustrate that consistent sequences had consistent object order across repetitions whereas random sequences did not.

#### 2.2.1. Learning session

The learning session included two study-test cycles. In each **study cycle**, the three consistent and the two random sequences were each presented three times. One of five semantic yes/no questions, e.g. “Is the presented item readily edible?”, was presented at the beginning of each cycle, and participants answered this question for each object presented within the cycle. Participants were instructed to answer as quickly and accurately as possible. The order of sequences was semi-randomised to ensure that sequences of the same type were not presented consecutively and that all sequences had been presented before showing repeats. Each object was displayed for 1 s and was followed by a blank fixation screen lasting 1.5 s. Participants could respond any time between object onset and the end of this fixation screen. The sequences were separated by a longer blank fixation screen lasting 2.5 s (see Fig. 2 for more detail).

**Figure 2.**
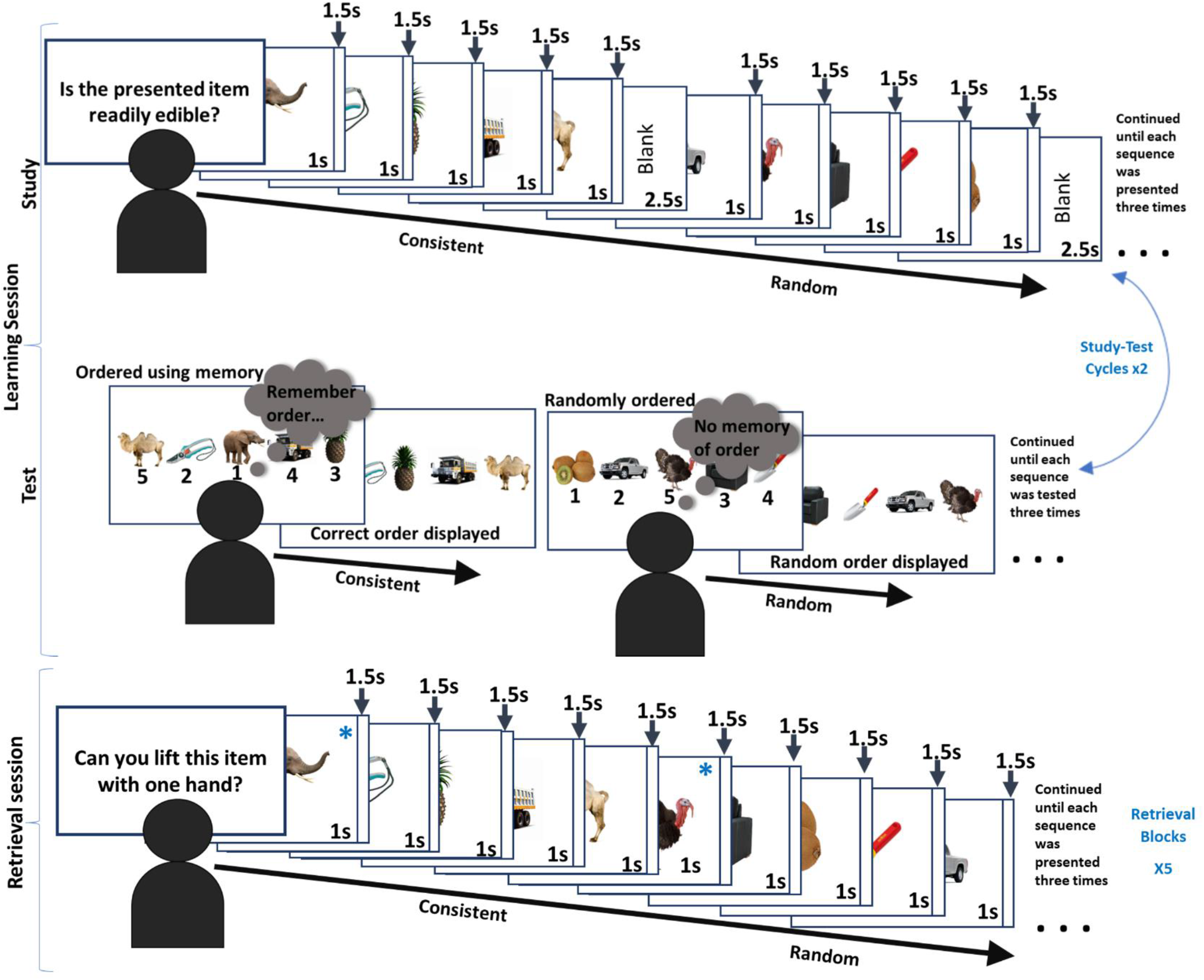
Sequence Memory Task Layout. The task comprised a “learning” session followed by a “retrieval” session. Examples of the consistent and random sequences are shown. The learning session comprised study-test cycles. In the study part, participants became familiar with the object sequences by answering semantic questions about the objects presented in their sequences. In the test part, participants were asked to re-order (now randomly ordered) objects from the consistently ordered sequences. Note that the random sequences cannot be reordered correctly, and participants randomly ordered the objects before being shown another random order. The sequences were shown three times within the test and study parts, and there were two study-test cycles. The retrieval session comprised five blocks, which was similar to the study part of the learning session, but all the sequences were shown seamlessly (without longer blank fixation screens). Again, participants answered semantic questions about each object displayed. Blue asterisks have been added for illustrative purposes, on some images in the retrieval session, to denote the image in position 1 of a sequence.

In each of the **test cycles** in the learning session, participants were shown all classes of sequences again. As above, the order of sequences was semi-randomised. Participants were explicitly tested on how well they had learned each of the consistent sequences, three times. They were shown all the objects from a sequence simultaneously and these were labelled 1 to 5, with five boxes underneath each of the numbers. Participants were asked to reconstruct the order in which they were presented in that sequence, using keys 1 to 5 along the top of the keyboard. The correct order was then displayed. For the random sequence trials, participants simply placed the objects in a random order and then another random order was displayed, which required no response. There were two study-test cycles within the learning session. For consistent sequences, answers were scored correct if objects were placed in the correct temporal position. The fraction of answers that were correct, within a sequence, was expressed as a percentage. Percentage accuracy over all the reconstruction tests was averaged to give a “Learning Session Performance”. Note that this score was comprised of explicit measures of sequence retrieval, assessed intermittently during the learning session, with the aim of reflecting learning performance.

#### 2.2.2. Retrieval session

Each of the five blocks was preceded by the presentation of a yes/no semantic question (different from those used in the learning session). Each sequence was presented three times within each block (again, in a semi-randomised order to ensure that sequences of the same type were not presented consecutively). The presentation times were the same as those in the learning session, except that the sequences were run seamlessly: sequence boundaries were not highlighted by a longer blank fixation screen.

RTs of positions 2-5 of the consistent and of the random sequences were averaged, and compared, to measure the extent to which individuals utilized temporal order knowledge to facilitate semantic judgments. This was expected to be reflected as a speeding in average response RT of the consistent sequences compared to the random sequences, “Sequence Memory Performance” (our measure of individual differences in sequence memory) (Crivelli-Decker et al., 2018; Hsieh et al., 2014). Note that RT of position 1 of each sequence was excluded as we expected equally slow RTs for position 1 across all sequences, as items in a sequence cannot be predicted until after the first item is presented.

### 2.3. MRI Data Acquisition

Structural and diffusion MRI data were collected using a Siemens Prisma 3T MRI system with a 32-channel head coil. Diffusion-weighted Imaging (DWI) data were acquired using a dual-shell HARDI (High Angular Resolution Diffusion-Weighted Imaging) (Tuch et al., 2002) protocol with the following parameters: slices = 80, Repetition Time (TR) = 9400 ms, Echo Time (TE) = 67 ms, Field of view (FOV) = 256 mm x 256 mm, acquisition matrix size 128 × 128, voxel dimensions = 2 × 2 × 2 mm^3^. Diffusion sensitization gradients were applied along 30 isotropic directions with a b-value of 1200 s/mm^2^ and along 60 isotropic directions with a b-value of 2400 s/mm^2^. Six non-diffusion-weighted images were also acquired with a b-value of 0 s/mm^2^.

T1-weighted anatomical images were obtained using a magnetization prepared rapid gradient echo scanning (MPRAGE) sequence with the following parameters: slices = 176, TR = 2250 ms, FOV = 256 mm x 256 mm, matrix size = 256 mm x 256 mm, flip angle = 9°, TE = 3.06 ms, slice thickness = 1 mm, voxel size: 1 × 1 × 1 mm^3^).

### 2.4. MRI Data Processing

The T1-weighted and DWI data were converted from DICOM to NIfTI formats using dcm2nii (obtained from www.nitrc.org). The T1-weighted data also underwent cropping, skull removal with the FSL Brain Extraction Tool (Smith, 2002), and down-sampling to a voxel size of 1.5 × 1.5 × 1.5 mm (Jenkinson et al., 2012).

Subject motion and echo planar imaging distortions were corrected by co-registering the DWI data to their respective T1-weighted images using Explore DTI (version 4.8.3) (Leemans, 2009). The lower (1200 s/mm^2^) and higher (2400 s/mm^2^) b-value data were analysed separately. Although DTI modelling was carried out on both shells, DTI model maps from the lower b-value data (where the assumption of Gaussian diffusion is met; Assaf & Pasternak, (2008)) were used to extract DTI scalar measures, FA (range 0 - 1) and Mean Diffusivity (MD, units 10^−3^ mm^2^/s), Radial Diffusivity (RD, units 10^−3^ mm^2^/s) and Axial Diffusivity (AD, units 10^−3^ mm^2^/s) (Leemans et al., 2009; MATLAB, 2015).

To estimate the diffusion tensor in the presence of physiological noise and system-related artefacts, the Robust Estimation of Tensors by Outlier Rejection algorithm was applied (Chang et al., 2005) to the lower b-value data. The ‘Free Water Elimination’ (FWE)-DTI two-compartment model technique (Pasternak et al., 2009) was used to allow removal of the free water contribution to the data, improving tissue specificity.

To detect and eliminate signal artefacts in the higher b-value data, the Robust Estimation in Spherical Deconvolution by Outlier Rejection algorithm was applied (Parker et al., 2012). Subsequently, peaks in the fibre Orientation Density Function (fODF) in each voxel were extracted using a modified damped Richardson-Lucy deconvolution algorithm (Dell’acqua et al., 2010). Whole-brain deterministic tractography was carried out in Explore DTI (version 4.8.3) (Leemans, 2009). The streamlines were constructed using an fODF amplitude threshold of 0.05, one seed per voxel, a step size of 0.5 mm and an angle threshold of 45°.

The dual-shell data was also processed to apply the biophysical NODDI model (Zhang et al., 2012), resulting in NDI (estimates the volume fraction of neurites) and ODI (estimates the variability of neurite orientation) values (both range 0 - 1), as well as a volume fraction of isotropic diffusion (Viso), attributed to a free water compartment. NODDI maps were created using Accelerated Microstructure Imaging via Convex Optimization (AMICO) (Daducci et al., 2015) NODDI (description and pipelines available here^1^). The resulting maps were accepted if normalized root-mean-square error map voxels in the areas of interest had error values under 0.3.

### 2.5. Extraction of Tract Streamlines

To generate three-dimensional streamlines that represented the fornix, the ILF and the PHC, ‘way-point’ ROIs were manually drawn onto whole-brain FA maps in the diffusion space of 18 subjects, using ExploreDTI (version 4.8.3) (Fig. 3). These ‘way-point’ ROIs allow the user to define Boolean AND and NOT gates, and SEEDS, with the aim of isolating the relevant streamlines. The protocol of Hodgetts et al. (2015) was used to extract fornix streamlines, with the exclusion of the AND gate placed on the transverse plane, as it did not appear to be required. The protocol of Wakana et al. (2007) was used to extract ILF streamlines. The protocol of Jones et al. (2013) was used to extract PHC streamlines, but a NOT gate placement of Sibilia et al. (2017) was used to exclude any streamlines of the cingulum that curved forward. The resultant tracts were used to train in-house automated tractography software (Greg Parker, Cardiff University; MATLAB, 2015), which was then applied to the entire dataset. Streamlines produced by the automated tractography software were visually inspected, and spurious fibres were removed using additional NOT gates.

**Figure 3.**
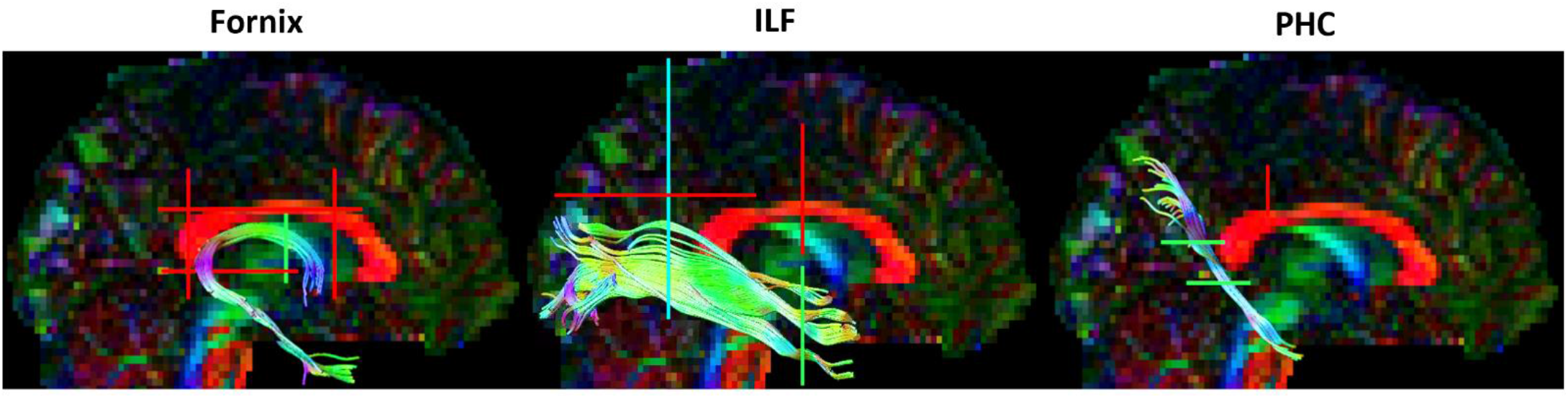
Construction of Tract Streamlines. Sagittal views of the fornix, ILF and PHC streamlines constructed in an ExploreDTI example dataset (available here: https://www.exploredti.com/exampledataset.htm). Right ILF and PHC tracts are shown, though they were extracted bilaterally. Colours on the brain map and the streamlines indicate diffusion along the gradient directions (left-right: red; top-bottom: blue; front-back: green). Example locations of the Boolean gates are represented by coloured lines (NOT gate: red. AND gate: green. SEED: blue). ILF: Inferior Longitudinal Fasciculus. PHC: Parahippocampal Cingulum.

### 2.6. Dimensionality Reduction of Microstructure Data

FA, MD, RD, AD (from FWE-DTI), and NDI and ODI (from NODDI) values for the voxels encompassed in the tract streamlines were extracted and averaged for each tract. This resulted in six microstructure metrics for three tracts for all 51 participants. These diffusion measures were reduced through PCA into microstructurally informative features (Chamberland et al., 2019; Geeraert et al., 2020; see also Gagnon et al., 2022; Guberman et al., 2022; Vaher et al., 2022). The tract microstructure data were combined in a single table. The Bartlett test was used to assess the appropriateness for PCA. The *prcmp* function in R (R Core Team, 2019) was then used to apply PCA to centred and scaled data (converted to z-scores). Sampling adequacy of the PCA results was tested using the Kaiser-Meyer-Olkin (KMO) test (from the R ‘Psych’ package; Revelle, 2022). Components were retained depending on the amount of cumulative variation they explained and on inspection of the scree plot. Following data reduction, participant scores in two biologically interpretable principal components (PCs) were used for analysis.

### 2.7. Statistical Analysis

For statistical testing and figure generation, R (R Core Team, 2019), RStudio (RStudio Team, 2020) and JASP (version 0.9.0.1) (JASP Team, 2021) were used. Participant datasets containing outlying values (>3 SDs from the mean) in either the behaviour condition or the microstructure PCA score data were identified and removed. T-tests to test differences between sequence condition RTs, and two-tailed Pearson’s tests for correlations between performance and microstructure scores were carried out in R. To correct for multiple comparisons, p-values were Bonferroni-corrected by dividing the standard 0.05 alpha level by the number of tracts: 0.017 (0.05/3 tracts).

In addition, Bayes Factors (BFs) were calculated using the BayesFactor package in R (Morey & Rouder, 2018), and reported as BF_10_ (evidence of the alternative over the null model). To aid interpretation, BF_10_ values between 1 and 3 were taken as weak evidence in favour of the alternative model, and values exceeding 3 were taken to reflect stronger evidence (Raftery, 1995). In contrast, values between 1 and 0.33, and below 0.33, were taken as weak and stronger evidence in favour of the null, respectively (Raftery, 1995). In the cases where behavioural scores needed to be controlled-for, Bayesian correlations between the residuals of variables were tested.

Plots were drawn using several R packages including ggcorrplot (Kassambara, 2019), ggstatsplot (Patil, 2021), ggplot2 (Wickham, 2016) and raincloudplots (Allen et al. 2021).

## 3.0. Results

### 3.1. Sequence Learning and Memory Data

#### 3.1.1. Learning session

Results from the reconstruction tests of the learning session showed that consistent sequences were learned reasonably well. The mean of the scores from the final repeat of the second cycle (the 6^th^ reconstruction of a sequence) was 90.39%. To characterize participant performance during the learning session, ‘Learning Session Performance’ scores were created by averaging the reconstruction results across Study-Test cycles. The mean (and SD) of Learning Session Performance was 80.82% (16.90%).

#### 3.1.2. Retrieval Session

The key retrieval session measure of interest, “Sequence Memory Performance” was calculated as the difference in averaged response RTs to positions 2-5 of the sequence types random and consistent. Figure 4A shows group RT data for each position in the two sequences. Figure 4B shows that average RTs to positions 2-5 were significantly faster for items in consistent sequences than for items in random sequences (t_(50)_ = 3.495, p = 0.001, d = 0.489).

**Figure 4.**
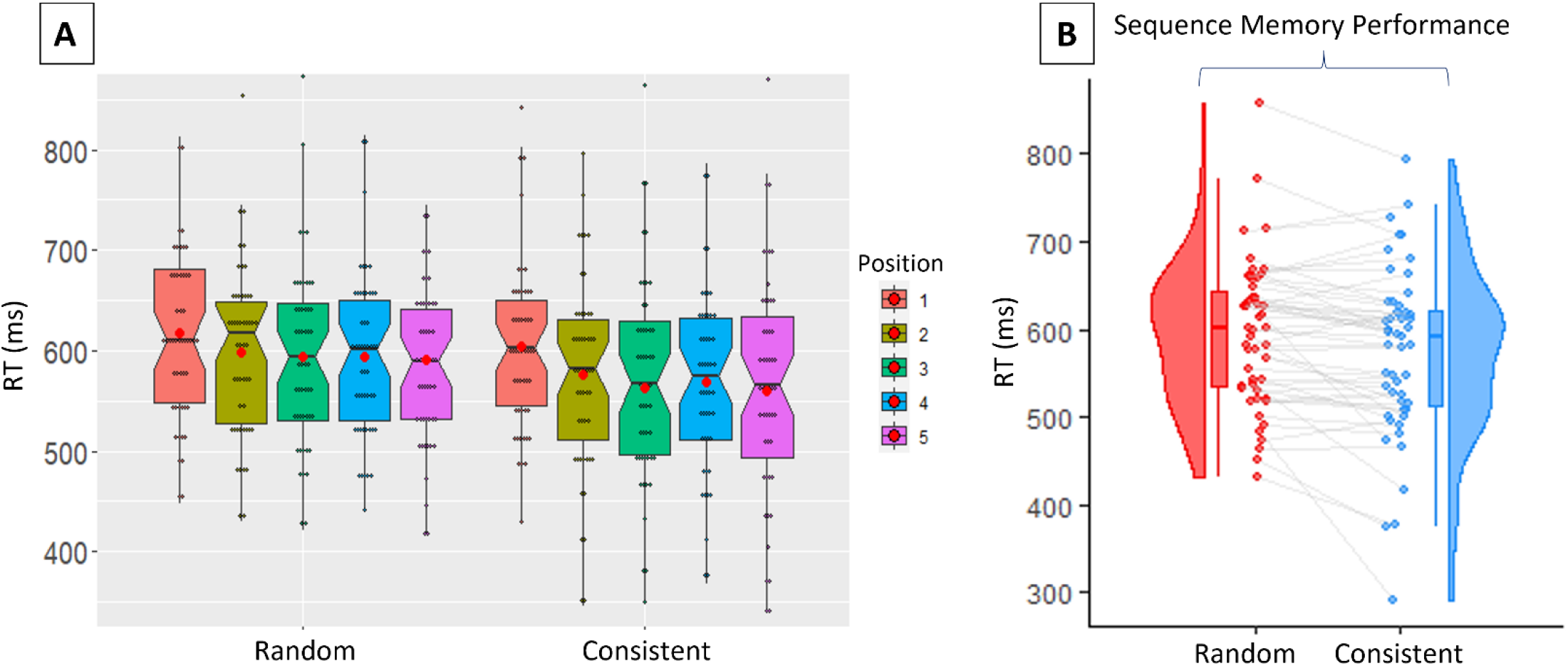
Participant Averaged Response RTs, for Each Condition. A) Box plots of the RTs for each position in the Random and Consistent sequence conditions. The red dots indicate the mean values. Sequence conditions are colour coded according to the key on the right. Note that the y-axis range was reduced to make differences in RTs between positions clearer, so some outlying high and low individual values (grey dots) have been excluded. B) Individual participant averaged response RTs, for positions 2-5, for the Random and Consistent sequence conditions. Data points from the same individuals are connected with grey lines. Sequence Memory Performance is the difference between random and consistent averaged RTs. Box plots and distributions are also displayed. Lines connected to the boxplots indicate 95% confidence intervals.

### 3.2. Tract microstructure

Fornix, ILF and PHC streamlines were successfully contructed in all participants. The means (and SDs) of the number of streamlines in each tract reconstruction were: fornix 546.6 (119.6); ILF 615.2 (183.0); PHC 119.1 (41.4). The means (and SDs) of the lengths in voxels of streamlines were: fornix 182.7 (11.6); ILF 203.7 (9.1); PHC 131.0 (14.9). Mean along-tract, bilaterally averaged tract microstructure metrics are shown in the *Supplementary Materials Table S1*. The Pearson correlation values shown in Figure 5A highlight the shared variance in these data.

**Figure 5.**
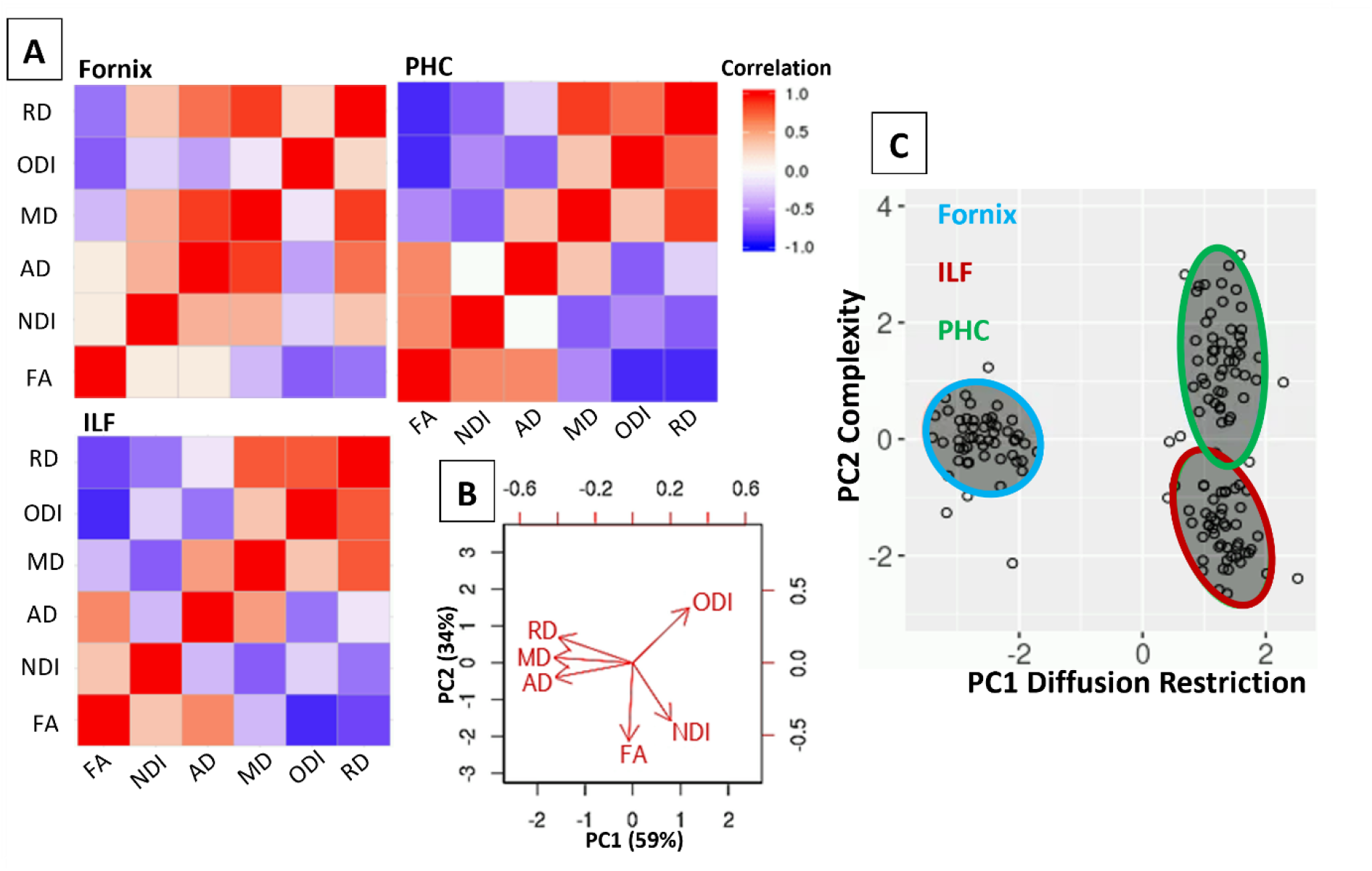
Redundancies Between Tract Microstructure Values and Reduction Through PCA. A) Pearson’s correlations within the microstructure data from each tract suggest that the DTI and NODDI metrics give overlapping information. Colour denotes r value. B) Biplot illustrating the influence of each of the measures on PC1 and PC2, which account for 59% and 34% of the variance, respectively. C) Tract component scores for each participant, illustrating the differing properties of the tracts. Note that one outlying PHC value lies beyond the boundaries of the graph. ILF: Inferior Longitudinal Fasciculus. PHC: Parahippocampal Cingulum. FA: Fractional Anisotropy. MD: Mean Diffusivity. NDI: Neurite Density Index. ODI: Orientation Dispersion Index. PC: Principal Component.

The results from the PCA (overall KMO: 0.63, sphericity: p<0.001; comparable to Chamberland et al., 2019 and Geeraert et al., 2020) showed that 93% of the microstructure data variability was accounted for by the first two principal components, PC1 and PC2 (Fig. 5; loadings shown in *Supplementary Materials Table S2*). PC1 accounted for 59% of the variance, and MD, RD and AD provided major negative contributions. PC1 resembled the first component found in Chamberland et al. (2019) and the second component of Geeraert et al. (2020) (see also Gagnon et al., 2022). Therefore, PC1 was interpreted as positively relating to a ‘diffusion restriction’ property of the fibres (presumably related to myelin density and axonal packing; henceforth referred to as ‘diffusion restriction’) (Beaulieu, 2014). PC2 accounted for 34% of the variance, and FA and NDI provided the major negative contributions, while ODI provided a major positive contribution. Since ODI is lower in tracts known to have greater fibre coherency and higher in tracts known to have more fibre fanning and crossing (Mollink et al., 2017; Zhang et al., 2012), and FA can be influenced by how coherently fibres within a voxel are organised (Jones, Knosche, et al., 2013; Pierpaoli et al., 1996), PC2 was interpreted as relating to a ‘complexity’ property of the fibres (the dispersion of modelled fibre orientations; henceforth referred to as ‘complexity’) (Schilling et al., 2018). Note that PC2 scores were sign-flipped, so that larger PC2 scores indicated greater complexity, to facilitate interpretation.

### 3.3. Relationships between sequence memory and tract microstructure

As is often the case with real/social science data (Bono et al., 2017), Sequence Memory Performance was non-normally distributed (right skew, >1) so, to allow the use of parametric testing throughout the analyses, the data were transformed. A constant (of the most negative value, sign-flipped and rounded up) was added to each value, and the square root was then calculated (McDonald, 2014), resulting in normally shaped data. Unless stated otherwise, ‘Sequence Memory Performance’ henceforth refers to the transformed data.

As the Extended Hippocampal System is known to be important for episodic memory (Gaffan & Hornak, 1997; Aggleton & Brown, 1999), we hypothesized that individual differences in fornix microstructure, which may reflect a quality of information flow between regions of this system (Jankowski et al., 2013), would relate to individual differences in sequence memory.

In line with our hypothesis, there was a significant positive correlation between Sequence Memory Performance and fornix microstructural complexity (such that better item-in-sequence memory was associated with higher complexity scores), and the resulting BF indicated evidence in favour of the alternative model (r_(46)_ = 0.343, p = 0.017, BF_10_ = 4.16). The correlation with fornix diffusion restriction, however, was not significant (r_(46)_ = 0.029, p = 0.847, BF_10_ = 0.33).

There were no significant correlations between Sequence Memory Performance and ILF diffusion restriction/complexity (r_(46)_ = 0.030, p = 0.839, BF_10_ = 0.33; r_(46)_ = 0.053, p = 0.720, BF_10_ = 0.34), or PHC diffusion restriction/complexity (r_(46)_ = −0.014, p = 0.923, BF_10_ = 0.33; r_(46)_ = 0.116, p = 0.433, BF_10_ = 0.43). These results indicate that microstructure of the fornix, specifically the ‘complexity’ component, relates to memory of objects in temporal context, whereas microstructure properties of the ILF and PHC do not.

Moreover, the correlation between Sequence Memory Performance and fornix complexity held when controlling for Learning Session Performance (r_(45)_ = 0.369, p = 0.011, BF_10_ = 6.11), indicating that this fornix property may be specifically important for sequence retrieval. There also continued to be no significant correlations between Sequence Memory Performance and fornix diffusion restriction, ILF diffusion restriction/complexity or PHC diffusion restriction/complexity, when controlling for Learning Session Performance (Fig. 6A).

**Figure 6.**
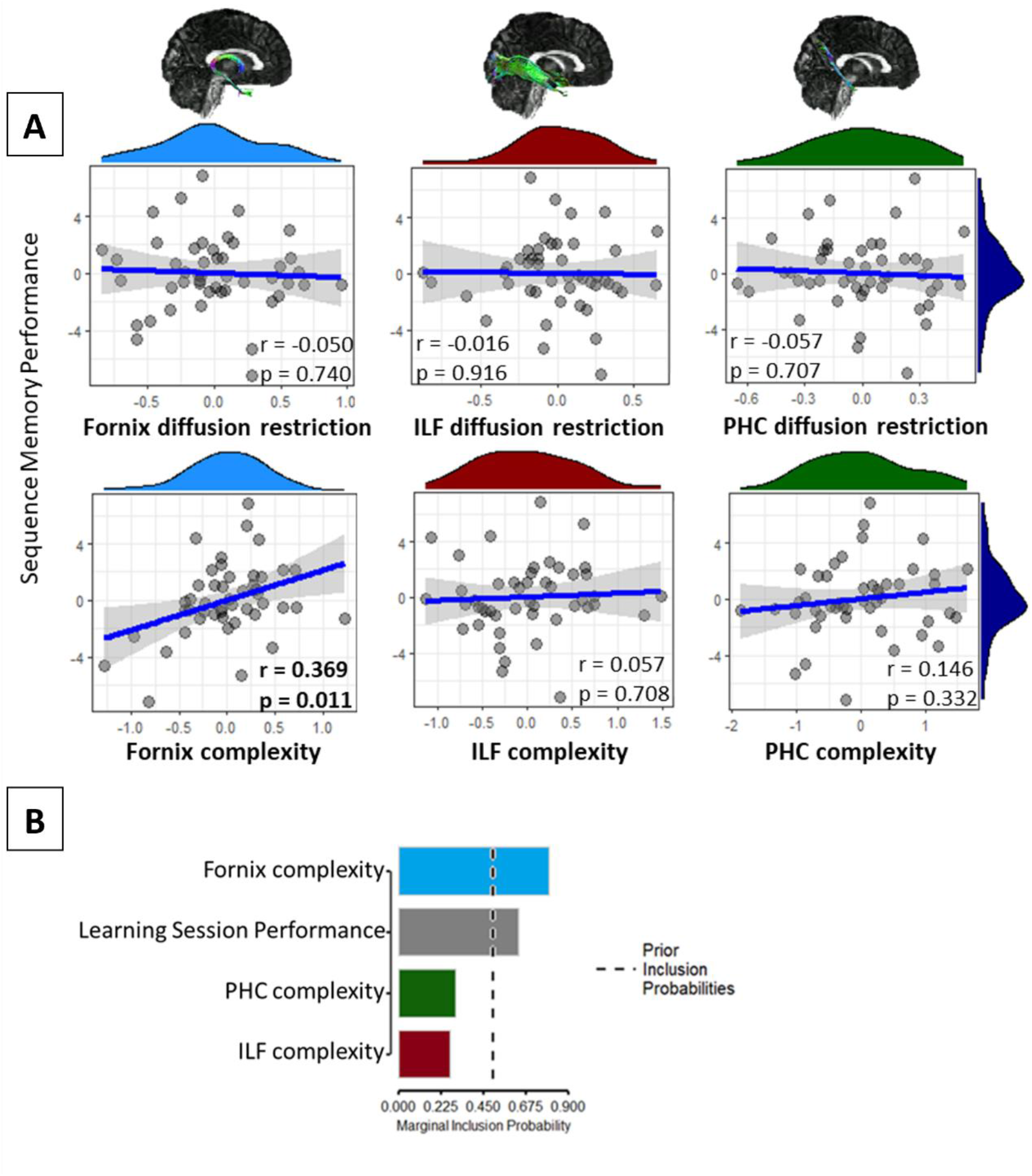
Associations Between Tract Microstructure Scores and Sequence Memory Performance. A) Correlation plots between Sequence Memory Performance and microstructure scores, when controlling for Learning Session Performance. The light-blue histograms show the distributions of the fornix diffusion restriction/complexity data. The green and red histograms show the distributions of the PHC diffusion restriction/complexity data and ILF diffusion restriction/complexity data, respectively. The dark blue histograms show the distribution of Sequence Memory Performance. The blue lines are the regression lines and surrounding shaded areas represent the 95% confidence interval. There was a significant partial correlation between Sequence Memory Performance and fornix complexity (at alpha threshold 0.017). Example fornix, ILF and PHC streamlines are shown on template brains. B) Bar graph of the posterior inclusion probabilities of the Bayesian linear regression, predicting Sequence Memory Performance with variables Fornix complexity, ILF complexity, PHC complexity and Learning Session Performance. The dashed line indicates the prior inclusion probabilities. ILF: Inferior Longitudinal Fasciculus. PHC: Parahippocampal Cingulum.

Bayesian multiple regression analysis, comparing models with fornix complexity, PHC complexity, ILF complexity and Learning Session Performance as predictors of Sequence Memory Performance (inclusion probabilities shown in Fig. 6B), indicated that a model with fornix complexity and Learning Session Performance best predicted Sequence Memory Performance (BF_10_ = 5.61, against the intercept-only model). This was followed by a model with only fornix complexity (BF_10_ = 3.05).

These findings indicate the relative importance of the fornix, compared with the PHC and ILF, especially the fibre complexity component of this tract, for memory of objects in temporal context.

## 4.0. Discussion

The hippocampus has been shown to play a crucial role in the construction and retrieval of representations of events associated through their temporal contiguity (Howard & Eichenbaum, 2015; Ranganath & Hsieh, 2016; Schapiro et al., 2016). Complementing this finding, here we demonstrate for the first time that microstructure of the fornix – a white matter tract that interconnects an extended hippocampal system, including the hippocampus, anterior thalamic nucleus, mamillary bodies and medial prefrontal cortex – predicts individual differences in temporal sequence memory in humans. These results lend credence to the idea that hippocampal contributions to memory may be supported by an extended network that is linked via the fornix (Aggleton, 2012; Aggleton & Brown, 1999; Bubb et al., 2017; Gaffan, 1992; Gaffan & Hornak, 1997).

Previously, through representational similarity analysis of fMRI signals recorded during the retrieval phase of a sequence memory task, Hsieh et al. (2014) demonstrated that individual differences in hippocampal voxel pattern information correlated with reaction time indices of sequence learning. Using scalp electroencephalography (EEG) recordings with the same task, Crivelli-Decker et al. (2018) showed that changes in oscillatory activity in the theta band (4-7 Hz) predicted sequence learning in this paradigm. Our findings complement these prior results in revealing that microstructure properties of the fornix, in particular a property reflecting fibre complexity, correlated with inter-individual RT difference between consistent and random sequences, confirming the importance of the hippocampus’s structural connectivity in supporting successful sequence memory.

Notably, in the Hsieh et al. (2014) paradigm, activity patterns in the parahippocampal cortex and perirhinal cortex carried information about temporal position and object identity respectively, but the hippocampus uniquely carried conjunctive representations of item and temporal position (see also Parker & Gaffan, 1996, 1997 for comparable evidence of a unique role for the primate fornix in integrating information about objects and their positions in space). Our results align with this by supporting a unique role of the fornix in item-in-temporal sequence memory. We found no evidence to suggest that microstructure properties of the ILF or PHC related to sequence memory, and the Bayes factors of these correlations indicated (weak) evidence in favour of the null. Moreover, supporting the unique role of the fornix, Bayesian multiple regression analysis, comparing models with fornix complexity, PHC complexity, ILF complexity and Learning Session Performance as predictors of Sequence Memory Performance indicated that a model with fornix complexity and Learning Session Performance best predicted Sequence Memory Performance. This preponderance of evidence supports the idea that retrieval of object sequences is supported specifically by an extended hippocampal system.

The fornix contains bidirectional pathways supporting direct hippocampal connections within the extended hippocampal system (Aggleton & Brown, 1999; Aggleton et al., 2010; Aggleton et al., 2015; Bubb et al., 2017). The fornix may contribute to hippocampal functioning by conveying theta rhythms, the prominent oscillation band of the hippocampus (Buzsaki, 2002), through this system (Aggleton et al., 2010; Jankowski et al., 2013). The fornix directly connects the hippocampus to: the septum/diagonal band of Broca, which supports the generation of theta activity (Leao et al., 2015; Swanson & Cowan, 1979); the supramammillary area, which has been shown to influence the frequency of hippocampal theta (Kirk, 1998; Pan & McNaughton, 2004); and the anterior thalamic nuclei, which not only operate in conjunction with the hippocampus through theta interactions, but also modulate hippocampal theta (Żakowski et al., 2017). The fornix may be both delivering theta rhythms to the hippocampus and facilitating hippocampal influence on extended hippocampal system theta. For example, fornix transection, disconnecting the septum from the hippocampus, abolishes hippocampal theta (Rawlins et al., 1979), while stimulating the fornical input from the hippocampus to anterior thalamic nuclei modulates thalamic theta (Tsanov et al., 2011; Jankowski et al., 2013).

Hippocampal theta is understood to be critical to the temporal organisation of active neuronal ensembles (Buzsaki, 2002), and may therefore support temporal organisation of episodic memory (Skaggs et al., 1996; Buzsáki and Moser, 2013; Herweg et al., 2020; Eichenbaum, 2017). Indeed, MTL theta phase coding is evident in humans learning image sequences (Reddy et al., 2021). Moreover, using electroencephalography in conjunction with a sequence memory task like that used by Hsieh et al., and in the current study, Crivelli-Decker et al. (2018) showed frontal midline theta power (which may be influenced by hippocampal theta, Hsieh & Ranganath 2014, through connections mediated by the fornix, Aggleton et al. 2015), to be lower during items in consistent sequences than during items in random sequences. Additionally, decreases in frontal midline theta power (which may reflect elevated theta connectivity, Solomon et al., 2019) correlated with RT of semantic decisions on upcoming objects in consistent sequences (Crivelli-Decker et al., 2018). It may be that hippocampus-influenced frontal midline theta aids coding of temporal information specifically, rather than temporal *and* spatial information, as temporal duration and not spatial distance has been shown to relate to frontal midline theta power modulation (Liang et al., 2021). Together, these results clearly link theta activity with retrieval of sequence knowledge. Therefore, sequence memory may rely on inter-regional functional communication within and beyond the extended hippocampal system, mediated by theta connectivity conveyed by the fornix, the latter in line with the findings from our study.

However, our interpretation, that the fornix supports sequence memory through its role in communication of theta-based processes between the hippocampus and an extended hippocampal system, is likely incomplete. One shortcoming is that the precise relationship between inter-regional theta synchronization and local activity is not currently well understood (Solomon et al., 2019). Furthermore, the fornix also carries some (albeit light) non-hippocampal connections, including those between the entorhinal cortex and the anterior thalamic nuclei (Saunders & Aggleton, 2007; Saunders et al., 2005), both of which have been shown to conduct forms of temporal coding (Bellmund et al., 2019; Nelson, 2021). Future studies could incorporate measurement of extended hippocampal theta rhythms into this study design and test for relationships between individual differences in hippocampal theta modulation, the extent of theta-based communication (e.g., phase/amplitude coupling), fornix microstructure and sequence retrieval performance. However, this may require invasive recording, as measuring electrophysiological signals from deep sources non-invasively with Magnetoencephalography, for example, is possible but it would be challenging to distinguish the individual regions of this extended hippocampal network (Pu et al., 2018).

In this study, extending previous work from our lab, that focused on DTI measures, we adopted a recently developed dimensionality reduction framework to take advantage of redundancies in dMRI measures (Chamberland et al., 2019; Geeraert et al., 2020, see also Guberman et al., 2022; Gagnon et al., 2022). By combining multiple measures through PCA (Chamberland et al., 2019; Geeraert et al., 2020), we found that the two major components, PC1 and PC2, were contributed to mostly by MD, RD and AD; and FA, NDI and ODI, respectively. We therefore considered these components to capture the properties of fibre diffusion restriction and complexity, respectively. The PCA components reported in this study share similarities with those reported in previous studies (Chamberland et al., 2019; Geeraert et al., 2020). PC1 is similar to the first component reported in Chamberland et al. (2019), which was also named ‘restriction’ and was negatively influenced by RD and positively influenced by a measure of fibre density. PC1 is also similar to the second component in Geeraert et al. (2020), named ‘myelin and axonal packing’ and influenced negatively by RD and MD and positively by NDI. PC2 overlaps with the first component in Geeraert et al. (2020), which they named ‘tissue complexity’ because it was influenced positively by FA and negatively by ODI. Although there are minor differences in the resulting components across studies (as would be expected when there are differences in the microstructure measures and tracts considered), the commonalities in the components confirm the usefulness of microstructure data reduction as a method to characterize tract properties.

The positive correlation between fornix PC2 and our reaction time indices of sequence memory suggests that increased fornix tissue complexity (reduced fornix axon coherence) relates to better sequence memory. The causes of inter-individual variation in white matter microstructure are not fully understood, but presumably reflect both environmental and genetic influences (Cahn et al., 2021; Luo et al., 2022). Studies examining white matter FA and ODI over age, indicate that tissue complexity changes during early development, perhaps supporting development of complex behaviours. For example, scores on the tissue complexity component in Geeraert et al. (2020) increased with age in children. Positive and negative correlations have also been identified in children between white-matter ODI and reading skill, and between FA and reading skill, respectively (Huber et al., 2019), suggesting increased tissue complexity is linked to better performance, even when controlling for age. Since our cohort comprises young adults, an age group in which age-related FA increases plateau and age-related ODI increases are slower than that of older adults (Chang et al., 2015), it is likely that we captured stable, peak inter-individual differences in fornix complexity. Related findings have linked increased white matter fibre complexity to better scene memory (Tavor et al., 2014) (see also Giacosa et al., 2016 for links between reduced fibre coherence and acquired expertise in adults). It may be that fornix complexity reflects individual differences in the extent to which fornix axons disperse to reach target sites (Mathiasen et al., 2019; Poletti & Creswell, 1977), although other factors, such as an increase in the presence of glial cells, could contribute (Gagnon et al., 2022).

A limitation with this study is that the sample used was entirely female, reflecting the availability of a participant sample for this work. We have no predictions about sex differences in temporal memory, so we anticipate that inclusion of male participants would result in similar findings, and consistent with this, previous work examining fornix microstructure and memory performance in young healthy adults reported no effects of participant sex (Hodgetts et al., 2020; see also Cahn et al., 2021). Considering our tractography methods, it should be noted that although the diffusion metrics used here have been histologically validated separately (Sato et al., 2017), components resulting from their reduction with PCA have not (Geeraert et al., 2020, Gagnon et al., 2022). It is not clear, therefore, that any one underlying tract property would be wholly reflected in one component (i.e., complexity could influence PC2, but it could also influence PC1 to a lesser extent). Furthermore, virtual tract renditions are created from estimations of water diffusion directionality, not from the anatomy itself, and characterization of fibres is limited by the MRI technique (e.g., the magnetic gradient amplitudes determine resolution thereby limiting the minimum detectable fibre diameter (McNab et al., 2013)). Although we constructed virtual tract renditions using anatomical knowledge and were informed by previous research, it is not possible to test the specificity of the tractography methods, for each participant, without knowing the true underlying anatomy (Schilling et al., 2020).

In summary, this study demonstrated the specific importance of the fornix in supporting memory of objects within a temporal context, extending previous work demonstrating the importance of the hippocampus as a key region supporting binding of spatial-temporal information (Ranganath, 2010; Graham et al., 2010). Furthermore, and going beyond previous DTI studies, use of a recently developed microstructure dimensionality reduction technique allowed us to posit that fornix fibre complexity may underlie inter-individual variation in sequence memory performance, presumably by mediating individual differences in network communication efficiency/complexity within an extended hippocampal-fornix system critical for episodic memory.

## CRediT author statement

Conceptualization: Katja Umla-Runge, Kim S Graham, Charan Ranganath

Resources: Katja Umla-Runge, Andrew Lawrence, Kim S Graham

Project administration: Katja Umla-Runge

Investigation: Katja Umla-Runge, Alison G. Costigan

Data Curation: Marie-Lucie Read

Formal Analysis: Marie-Lucie Read

Methodology: Liang-Tien Hsieh, Maxime Chamberland

Writing: all authors

Supervision: Katja Umla-Runge, Kim S Graham, Andrew Lawrence, Charan Ranganath

## Data Availability Statement

Anonymised output data that support the findings of this study are available from the corresponding author upon reasonable request.

## Acknowledgments

We would also like to thank Matt Jones and Angharad Williams for their help with initial data curation, Ashvanti Valji and Vera Dehmelt for their help with data collection, and Greg Parker for the use of his Automated Tractography Software.

## Graphical Abstract

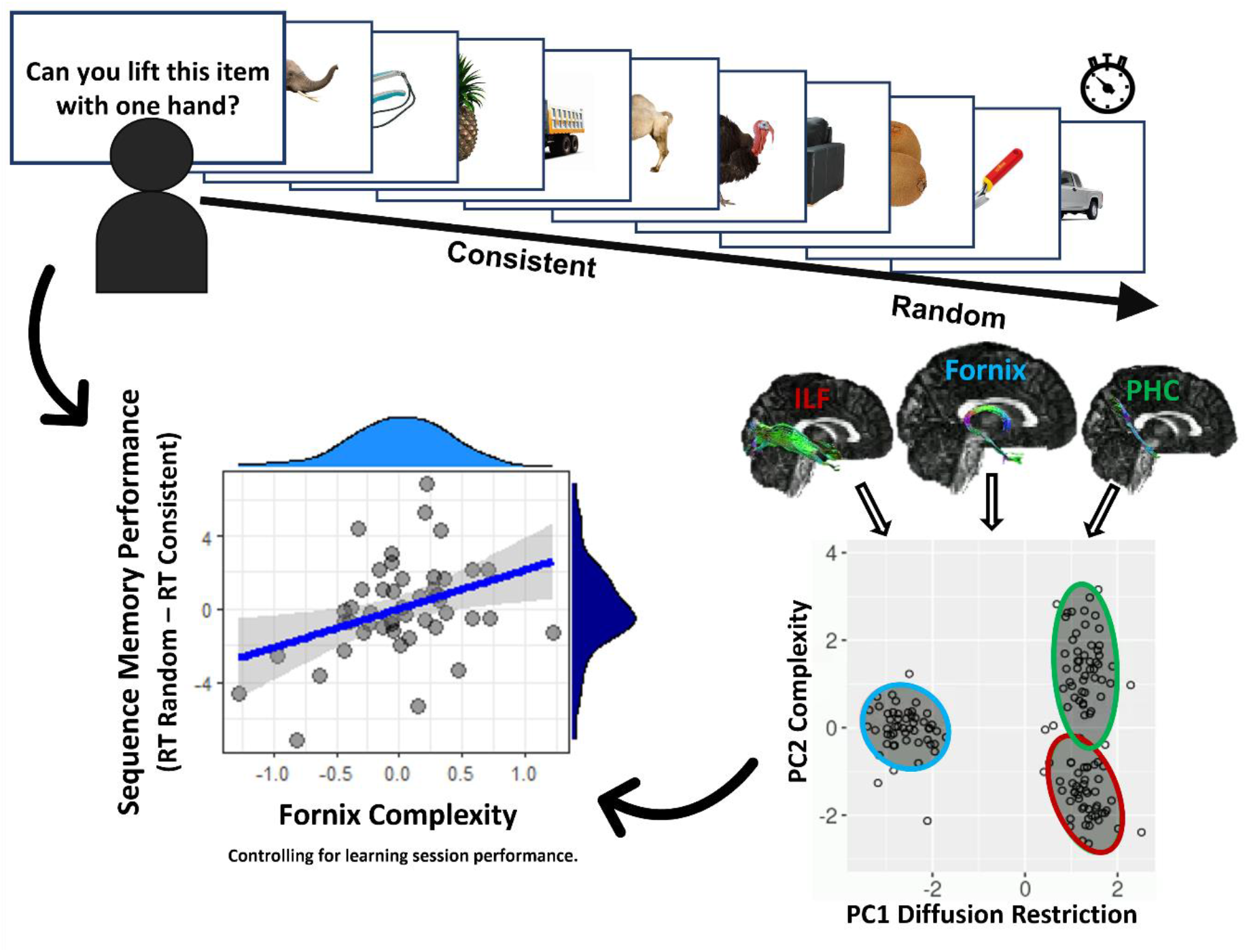

We tested whether individual differences in fornix microstructure, the main white-matter tract connecting the hippocampus, predicted sequence memory, by measuring participants’ differences in Reaction Times (RT) to answer semantic questions on objects presented in learnt consistently-ordered sequences and in randomly-ordered sequences (Sequence Memory Performance). Microstructure properties of the fornix, and two tracts not predominantly connecting the hippocampus: the Parahippocampal Cingulum bundle (PHC) and the Inferior Longitudinal Fasciculus (ILF), were reduced, using principal components analysis, into ‘fibre restriction’ and ‘complexity’. Fornix, but not PHC and ILF, complexity correlated with Sequence Memory Performance, highlighting a role for the fornix in temporal sequence memory.

## Footnotes

1 https://github.com/daducci/AMICO/blob/master/doc/demos/NODDI_01.md

