## Supplementary Materials for "A Role for the Fornix in Temporal Sequence Memory"

| | **Fornix** | | **ILF** | | PHC | |
|---|---|---|---|---|---|---|
| | **Group mean** | **SD** | **Group mean** | **SD** | **Group mean** | **SD** |
| **FA** | 0.40 | 0.01 | 0.44 | 0.02 | 0.35 | 0.03 |
| **MD** | 0.09 $x10^{-2}$ | 0.03 $x10^{-3}$ | 0.07 $x10^{-2}$ | 0.01 $x10^{-3}$ | 0.07 $x10^{-2}$ | 0.01 $x10^{-3}$ |
| **RD** | 0.07 $x10^{-2}$ | 0.03 $x10^{-3}$ | 0.05 $x10^{-2}$ | 0.02 $x10^{-3}$ | 0.06 $x10^{-2}$ | 0.02 $x10^{-3}$ |
| **AD** | 0.14 $x10^{-2}$ | 0.05 $x10^{-3}$ | 0.11 $x10^{-2}$ | 0.03 $x10^{-3}$ | 0.10 $x10^{-2}$ | 0.02 $x10^{-3}$ |
| **NDI** | 0.45 | 0.03 | 0.51 | 0.03 | 0.46 | 0.02 |
| **ODI** | 0.15 | 0.01 | 0.19 | 0.02 | 0.24 | 0.02 |

**Table S1. Mean along-tract, bilaterally averaged tract microstructure metrics.**

| | **PC1** | **PC2** |
|---|---|---|
| **FA** | -0.027 | -0.674 |
| **MD** | -0.527 | 0.046 |
| **AD** | -0.521 | -0.126 |
| **RD** | -0.495 | 0.222 |
| **NDI** | -0.255 | -0.502 |
| **ODI** | 0.375 | 0.476 |

**Table S2. PCA loadings.**
